# Multi-omic tumor data reveal diversity of molecular mechanisms that correlate with survival

**DOI:** 10.1101/267245

**Authors:** Daniele Ramazzotti, Avantika Lal, Bo Wang, Serafim Batzoglou, Arend Sidow

**Affiliations:** Department of Pathology, Stanford University, CA 94305, USA; Department of Computer Science, Stanford University, CA 94305, USA; Department of Genetics, Stanford University, CA 94305, USA

## Abstract

Outcomes for cancer patients vary greatly even within the same tumor type, and characterization of molecular subtypes of cancer holds important promise for improving prognosis and personalized treatment. This promise has motivated recent efforts to produce large amounts of multidimensional genomic (‘multi-omic’) data, but current algorithms still face challenges in the integrated analysis of such data. Here we present Cancer Integration via Multikernel Learning (CIMLR), a new cancer subtyping method that integrates multi-omic data to reveal molecular subtypes of cancer. We apply CIMLR to multi-omic data from 36 cancer types and show significant improvements in both computational efficiency and ability to extract biologically meaningful cancer subtypes. The discovered subtypes exhibit significant differences in patient survival for 27 of 36 cancer types. Our analysis reveals integrated patterns of gene expression, methylation, point mutations and copy number changes in multiple cancers and highlights patterns specifically associated with poor patient outcomes.

## Introduction

Cancer is a heterogeneous disease that evolves through many pathways, involving changes in the activity of multiple oncogenes and tumor suppressor genes. The basis for such changes is the vast number and diversity of somatic alterations that produce complex molecular and cellular phenotypes, ultimately influencing each individual tumor behavior and response to treatment. Due to the diversity of mutations and molecular mechanisms, outcomes vary greatly and it is therefore important to identify cancer subtypes based on common molecular features, and then correlate those with outcomes. This will lead to an improved understanding of the pathways by which cancer commonly evolves, as well as better prognosis and personalized treatment.

Efforts to distinguish subtypes are complicated by the many kinds of genomic changes that contribute to cancer - for example, point mutations, DNA copy number aberrations, DNA methylation, gene expression, protein levels, and post-translational modifications. While gene expression clustering is often used to discover subtypes (e.g., the *PAM50* subtypes^1^ of breast cancer), analysis of a single data type does not typically capture the full complexity of a tumor genome and its molecular phenotypes. For example, a copy number change may be biologically relevant only if it causes a gene expression change; gene expression data ignores point mutations that may alter the function of the gene product; and point mutations in two different genes may have the same downstream effect, which may become apparent only when also considering methylation or gene expression. Therefore, comprehensive molecular subtyping requires integration of multiple data types, which is now possible in principle thanks to projects such as The Cancer Genome Atlas (TCGA) and the International Cancer Genome Consortium (ICGC) that have generated multi-omic data on thousands of tumors.

In order to use multiple data types for subtyping, some approaches carry out separate clustering of each data type followed by manual integration of the clusters^2^. However, clusters based on different data may not be clearly correlated. More rigorous methods for integration include pathway analysis on multi-omic data, followed by clustering on the inferred pathway activities^3^, constructing fused networks of similarities between tumors^4^, rank matrix factorization^5^ and Bayesian consensus clustering^6^. There are also several sparse clustering methods, which assume that only a small fraction of features are relevant; for example, iCluster+^7^ uses generalized linear regression with lasso penalty terms. These methods are either highly dependent on preliminary feature selection, or enforce sparsity, thus neglecting potentially useful information. A recent method, PINS^8^, introduces a novel strategy of identifying clusters that are stable in response to repeated perturbation of the data.

One drawback common to many of the more principled methods is that they are computationally too intensive to be routinely applied to large data sets, due to the need for either parameter selection or repeated perturbations. Moreover, they treat all data types equally, which may not be biologically appropriate. As a result, in many cases, the discovered clusters show poor association with patient outcomes^9,10^. We therefore set out to develop a novel method that does not have these drawbacks.

CIMLR is based on SIMLR, an algorithm for analysis of single-cell RNA-Seq data^11^. CIMLR learns a measure of similarity between each pair of samples in a multi-omic dataset by combining multiple gaussian kernels per data type, corresponding to different but complementary representations of the data. It enforces a block structure in the resulting similarity matrix, which is then used for dimension reduction and k-means clustering. CIMLR is capable of incorporating complete genomes and of scaling to a large number of data types, and also does not assume equal importance for each data type. As such, it is well suited to modeling the heterogeneity of cancer data.

Here we apply CIMLR to discover integrative subtypes within 36 types of cancer. We recover known as well as novel subtypes, and show that our method outperforms current state-of-the-art tools in speed, accuracy, and prediction of patient survival. Moreover, for 5 of these cancers, we also validated the significant survival associations we discovered on unseen external datasets. This systematic subtype analysis, the most comprehensive to date, provides valuable insights into the biology underlying tumor variability.

## Results

### Subtyping of 36 cancer types using CIMLR

We carried out a systematic, integrative subtype analysis using CIMLR (Fig. 1a) across all 32 cancer types available from TCGA on a total of 6645 patients. Four data types were considered: point mutations, copy number alterations, promoter CpG island methylation and gene expression.

**Figure 1.**
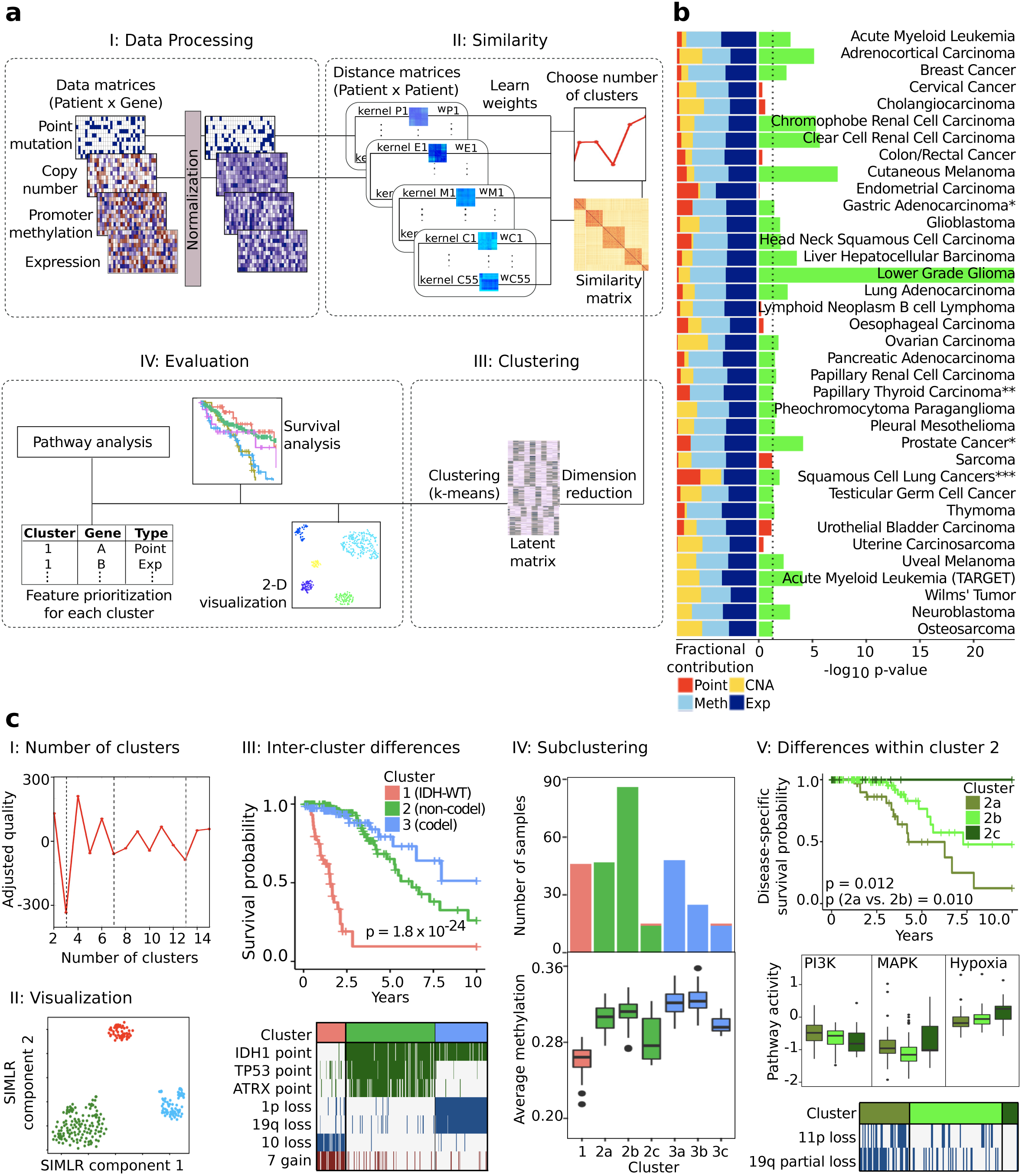
**(a)** CIMLR workflow. I: Each data type is arranged as a matrix where rows are patients and columns are genes. All matrices are then normalized so that values range from 0 to 1, so that all data types have the same range. II: For each data type, CIMLR learns weights for multiple kernels (each kernel is a measure of patient-to-patient distance). The number of clusters C is determined by a heuristic based on the gap statistic. The method then combines the multiple kernels into a symmetric similarity matrix with C blocks, where each block is a set of patients highly similar to each other. III: The learned similarity matrix is then used for dimension reduction and clustering into subtypes. IV: The clusters are evaluated by visualization as a 2-D scatter plot and survival analysis. The molecular features significantly enriched in each cluster are listed, and finally, pathway activity is compared. **(b)** Left: Contributions (measured as fraction of total kernel weight) by each data type. Right: Results of survival analysis on the best clusters for 32 cancer types. Green bars represent the 27 cancer types for which significant differences in patient survival were obtained between clusters; red bars represent the remaining cancers. For prostate cancer (*), significance for disease-free survival is shown as nearly all patients survived for the duration of the study. **(c)** Validation of our method on 282 lower-grade gliomas. I: Plot of separation cost showing 3 as the best number of clusters and 7 and 13 as secondary peaks. II: Separation between clusters. III: Differences in survival (upper) and molecular features (lower) between clusters. IV: Further separation into 7 subclusters (upper) with different methylation (lower; y-axis shows average beta value per patient). V: Differences in survival (upper), pathway activity (middler) and molecular features (lower) between the three subclusters of cluster 2. *PFI; **DSS; ***DFI. Otherwise: overall survival.

We evaluated the clusters produced by CIMLR based on (1) survival analysis, (2) silhouette (a measure of cohesion and separation of clusters^12^), (3) stability of the clusters, and (4) significant differences in pathway activity between clusters. Details of these calculations are given in the Methods and all values are given in Supplementary Tables 1-3. To demonstrate the value of multi-omic subtyping, we compared the performance of CIMLR using all 4 data types against analysis using only methylation or expression. We also compared CIMLR to four state-of-the-art existing methods for integrative subtyping: iCluster+^7^, Bayesian consensus clustering^6^, PINS^8^, and SNF^4^. CIMLR outperformed all other methods on all tested metrics. In particular, the clusters obtained using CIMLR show significant differences in patient survival in 23 of 32 cancer types from TCGA (Fig. 1c, Supplementary Table 1), exceeding the performance of all other approaches.

Additionally, we applied CIMLR to discover subtypes of pediatric cancers using data from the TARGET initiative^13^. In this case we considered 4 tumor types: Acute Myeloid Leukemia, Wilms tumor, Neuroblastoma and Osteosarcoma. Remarkably, the clusters obtained by CIMLR present significant differences in overall survival for all four pediatric tumor types (Table 1; Supplementary Tables 5-8).

**Table 1:** Comparison of subtypes discovered using CIMLR with multi-omic data, CIMLR with methylation or expression only, and other integrative subtyping methods, across 36 cancer types. Overall survival: Number of cancer types for which a significant (p<0.05) difference in overall survival was observed between clusters. All survival metrics: Number of cancer types for which a significant (p<0.05) difference in any available survival metric was observed between clusters. Stability: Normalized Mutual Information (NMI) of the clusters discovered by repeated k-means clustering (for methods that use k-means clustering). Pathway activity: Number of cancer types for which a significant (p<0.05) difference in the activity of any of 11 cancer-associated signaling pathways was observed between clusters. Data for pathway activity was available for 27 cancer types^17^. See Methods for details of calculations. Silhouette and stability are reported as mean +/- standard deviation.

|  | **CIMLR**<br>**(multi-omic)** | CIMLR<br>(methylation<br>only) | CIMLR<br>(expression<br>only) | iCluster+<sup>7</sup> | BayesCC<sup>6</sup> | PINS<sup>8</sup> | SNF<sup>4</sup> |
| --- | --- | --- | --- | --- | --- | --- | --- |
| TCGA (32 adult cancers, 4 data types) |  |  |  |  |  |  |  |
| Overall survival | **19** | 12 | 12 | 14 | 10 | 12 | 12 |
| All survival metrics | **23** | 14 | 20 | 18 | 11 | 17 | 16 |
| Silhouette | **0.81+/-0.22** | 0.81+/-0.09 | 0.84+/-0.09 | -0.07+/-0.54 | 0.09+/-0.08 | 0.00+/-0.07 | 0.00+/-0.00 |
| Stability | **0.89+/-0.08** | 0.91+/-0.05 | 0.91+/-0.04 | 0.25+/-0.32 | NA | NA | NA |
| Pathway activity | **26** | 24 | 26 | 17 | 21 | 26 | 25 |
| TARGET<sup>13</sup> (4 pediatric cancers, 3 data types) |  |  |  |  |  |  |  |
| Overall survival | **4** | 2 | 3 | 1 | 0 | 1 | 3 |
| Silhouette | **0.86+/-0.05** | 0.83+/-0.08 | 0.84+/-0.09 | 0.26+/-0.16 | 0.09+/-0.11 | 0.06+/-0.10 | -0.20+/-0.03 |
| Stability | **0.94+/-0.04** | 0.95+/-0.06 | 0.93+/-0.05 | 0.68+/-0.14 | NA | NA | NA |
| TCGA + TARGET (36 cancers) |  |  |  |  |  |  |  |
| All survival metrics | **27** | 16 | 23 | 19 | 11 | 18 | 19 |

One improvement provided by CIMLR over previous methods is that it learns weights for each data type instead of assigning equal importance to each. The contributions of each data type, measured as the fraction of total kernel weight contributed by kernels based on that data type, are very different between cancers (Figure 1b). Expression and methylation each contribute 30-50% of the kernel weight in almost all cancers; however, squamous cell lung cancers have a very low kernel weight for methylation. On the other hand, the contribution of point mutations and copy number are highly variable. We observe some association between these kernel weights and the C/M classification of cancer types^14^, with M-type cancers such as endometrial and colorectal cancers having high contributions from point mutations while copy number changes present higher contributions to subtyping of some C-type cancers such as ovarian cancer. CIMLR can thus give us insight into which data types are most useful and informative for subtyping in different cancers.

Finally, all other approaches except SNF proved impractically time-consuming and computationally intensive to run (on the order of days using 64 cores for a single configuration), while CIMLR takes minutes to run on a laptop for each cancer type. In summary, we find that multi-omic data integration using CIMLR is the most effective method for integrative subtyping based on technical performance, discovery of clinical and biological differences, and practical usability.

### Biological validation of CIMLR on lower-grade gliomas

Lower-grade (also called low-grade) gliomas are a well-studied example for genomic subtyping, which is why we chose it for validation of CIMLR via reproduction of robust, known results. Three subtypes of lower-grade gliomas have been characterized^15^, based on the presence of IDH1/2 point mutations and chromosome 1p/19q codeletion.

CIMLR reproduces the known subtypes of lower-grade gliomas (Fig. 1c), with 3 being the best number of clusters and additional peaks at 7 and 13 clusters. The 3 clusters found by CIMLR show strong separation and correspond to the known molecular subtypes. Cluster 1 is composed almost entirely of IDH-wild type samples with a characteristic loss of chromosome 10 and gain of chromosome 7. Cluster 2 (non-codel) is composed of mostly IDH mutant samples with additional point mutations in TP53 and ATRX. Cluster 3 (codel) is composed of IDH mutant tumors with a chromosome 1p/19q codeletion. The IDH-wild type cluster has the worst overall and disease-specific survival, followed by the non-codel cluster, while the codel group has the best outcomes.

A recent study^2^ comprising lower-grade gliomas and glioblastomas hinted at a possible finer classification of these tumors, finding a “CIMP-low” subgroup of IDH mutant non-codel tumors, with lower methylation and worse survival than the rest of the non-codel group. The codel group, on the other hand, was not divided further. In order to further characterize lower-grade gliomas, we investigated the results by CIMLR for 7 clusters, which are near-perfect subsets of the 3 major clusters. We find that in these results the codel and non-codel groups are divided into 3 subclusters each. In both groups, there are two CIMP-high subclusters and one CIMP-low subcluster.

We examined the subclusters of cluster 2. Subcluster 2c is characterized by reduced methylation, similar to the CIMP-low subgroup described previously^2^, while 2a and 2b have higher methylation levels. The three subclusters have significantly different overall (p=0.043) and disease-specific (p=0.012) survival. Analysis of gene expression patterns using PROGENy showed that the three subclusters also have significantly different activity of the PI3K, MAPK, and hypoxia pathways. Further, subcluster 2a, which has the worst survival outcomes, is associated with more copy number changes than 2b or 2c; 68% of samples in 2a have a partial loss of 19q (19q13.31-13.43), unlike the complete-arm loss in the codel group. 57% have a loss of 11p, including the tumor suppressor TRIM3, which also showed reduced expression in the same samples. Loss of TRIM3 has been associated with increased proliferation and stem cell-like properties of glioblastomas^16^.

Thus, CIMLR reproduces known molecular subtypes and also reveals novel subgroups within lower-grade gliomas. This provides empirical evidence that CIMLR can discover meaningful and robust biological subtypes on the basis of multi-omic data.

As for lower-grade gliomas, we evaluated the clusters found by CIMLR for all the cancer types on the basis of cluster separation and survival analysis. We see that CIMLR exceeds the performance of all other methods in identifying clusters associated with significant survival differences. While lower-grade gliomas were separated into 3 clusters with the most significant survival difference (p=1.8×10^-24^), we also obtained strong survival differences for cancers that have proven much more difficult to subtype, such as clear cell renal cell carcinoma (p=1.9×10^-6^).

To characterize the biological changes that lead to survival differences between clusters, we selected genetic alterations that were enriched in specific clusters, and used GSEA (Gene Set Enrichment Analysis) and PROGENy^17^ to identify cancer-related biological pathways that were activated differently between clusters. We then considered each individual cancer type and searched for features that might be related to the observed differences in survival. Below we present results for 8 selected cancers where we obtain a significant difference in survival and improve over previous clustering studies.

### Liver hepatocellular carcinoma

Hepatocellular carcinoma is associated with several risk factors including chronic hepatitis B virus (HBV) and hepatitis C virus (HCV) infection, and alcohol consumption. iCluster+ has been used to find 3 integrative subtypes^18^; however, there was no significant difference in patient outcomes, although some differences were seen in an external cohort that was tracked over a longer time. CIMLR finds 8 clusters (Figure 2a), associated with significant differences in overall and disease-specific survival within the cohort.

**Figure 2.**
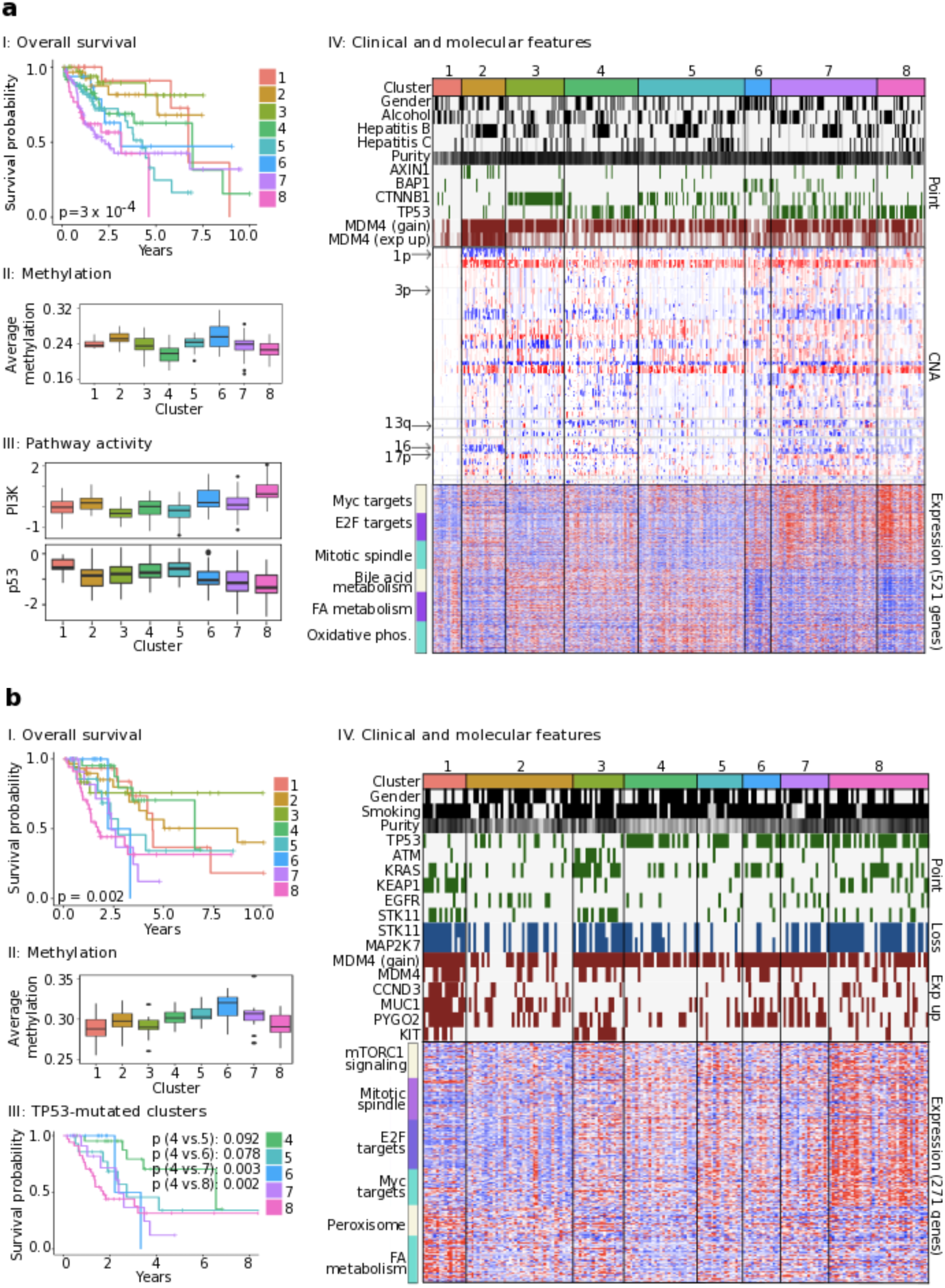
**(a)** Subtyping of 359 liver hepatocellular carcinomas. I: Overall survival probability for the 8 clusters. X-axis denotes years from diagnosis. II: Boxplots showing average promoter methylation (beta value) per patient, for each cluster. III: Boxplot showing PROGENy pathway activities for PI3K (upper) and p53 (lower) pathways, for each cluster. IV. Selected clinical and molecular features that differentiate the 8 clusters. For gender, alcohol, Hepatitis B, and Hepatitis C, gray bars represent missing data. Black bars represent females, alcohol consumption, Hepatitis B or Hepatitis C infection. Copy number alterations (CNA) and RNA expression are shown along a blue (less) to red (more) spectrum. **(b)** Clustering of 188 lung adenocarcinomas. I: Overall survival probability for the 8 clusters. II: Boxplots showing average promoter methylation (beta value) per patient, for each cluster. III: Overall survival probability for the 5 clusters associated with TP53 mutations. IV: Selected clinical and molecular features that differentiate the 8 clusters. For gender and smoking, gray bars represent missing data. Black bars represent females and tobacco smokers respectively.

Clusters 1, 2, and 3 have relatively high overall and disease-specific survival. We do not observe any common point mutations or copy number changes in cluster 1; however, this may be due to the low purity and higher immune infiltration of these tumors^19^. Cluster 2 also has high overall and disease-specific survival, and is associated with HBV infection (60% samples) and Asian ethnicity. Although most of these tumors are wild-type for TP53, they show gain and increased expression of the p53 repressor MDM4, and low p53 activity according to PROGENy. This cluster also has a universal loss on chromosome 1p including the succinate dehydrogenase gene SDHB, accompanied by reduced SDHB expression. Reduced SDHB blocks respiration leading to a metabolic shift toward glycolysis; the accumulation of succinate also inhibits demethylases leading to a CIMP (high methylation) phenotype^20^, which we in fact observe in this cluster. This cluster also displays losses on chromosome 16; this includes the tumor suppressors CYLD and TSC2, and the DNA repair gene PALB2, as well as reduced expression of all three. It is also enriched (28% samples) for mutations in AXIN1, a tumor suppressor gene that regulates the Wnt signaling pathway. GSEA shows that this cluster is enriched for tumors with reduced expression of genes for oxidative phosphorylation and the G1/S checkpoint.

Cluster 3 is enriched for mutations in CTNNB1 (beta-catenin). While CTNNB1 mutations are also common in other clusters, the tumors in cluster 3 also display high expression of GLUL (Glutamine synthase), a well-characterized target of beta-catenin^21^, suggesting that beta-catenin activation leads to glutamine synthesis and cellular proliferation in these tumors.

Patients in cluster 6 are more likely to be female (p=0.001), non-drinkers, and do not have HBV or HCV infection. This cluster is enriched for mutations in the histone deubiquitinating tumor suppressor BAP1, which is involved in chromatin remodeling as well as double-strand break repair (42% samples). 63% of samples also share a loss of BAP1 on 3p, along with reduced expression. These tumors have high DNA methylation, a phenotype previously associated with BAP1 mutations in renal cancers^22^, and frequently lack the 8p loss/8q gain that is seen in the other clusters. In addition, they show strongly reduced expression of genes for normal hepatocyte functions such as bile acid metabolism, fatty acid metabolism, xenobiotic metabolism, and coagulation.

Clusters 4, 7, and 8 are associated with TP53 point mutations as well as losses on 13q (RB1) and 17p (MAP2K4, TP53). While all of them also show reduced expression of RB1 and MAP2K4, Cluster 4 has stronger reduction in TP53 expression. However, clusters 7 and 8 have significantly worse overall survival than cluster 4 (p=0.045 and p=0.036 respectively). Both show increased expression of Myc and E2F target genes as well as genes involved in mTORC1 signaling and the mitotic spindle. In addition, cluster 8 shows reduced expression of genes involved in normal hepatocyte function (as seen in cluster 6), higher immune infiltration and macrovascular invasion. PROGENy scores show that p53 and PI3K pathway activities are significantly associated with the clusters (p < 10^-12^ for both), with cluster 8 showing the lowest p53 activity and highest PI3K activity.

### Lung adenocarcinoma

Lung adenocarcinoma, often caused by smoking, is the leading cause of cancer death globally. Previous studies identified transcriptional^23^ and histological^24^ subtypes, as well as 6 integrated clusters^9^, which, however, showed no significant association with patient survival. We find 8 clusters, significantly associated with overall and disease-specific survival (Figure 2b).

Clusters 1-3 are predominantly wild-type for TP53, whereas the remaining clusters (4-8) are associated with TP53 mutations. In general, the TP53 mutant clusters (4-8) are associated with worse survival outcomes; however, the exception is cluster 4, which has significantly better overall and disease-specific survival outcomes than the other TP53-mutant clusters, comparable to clusters 1-3.

Cluster 1 is characterized by loss of 19p, including the tumor suppressor STK11; this is associated with reduced STK11 expression. It is enriched for point mutations in STK11 and KEAP1, as well as high expression of CCND3 (cyclin D3), the transcriptional regulator MUC1, the Wnt pathway activator PYGO2 and the p53 inhibitor MDM4. In addition, it shows low DNA methylation, high expression of genes for fatty acid metabolism and peroxisome function, and low expression of genes involved in apoptosis and the G2/M checkpoint.

Cluster 3, like cluster 1, has low methylation, and is associated with STK11 loss and point mutations. In addition, it is enriched for point mutations in ATM and KRAS. It has a gain on 14q and losses on 1p, 21q (BTG3, PRMT2, HMGN1), and 15q (FAN1), as well as reduced expression of those genes. This cluster is associated with high expression of the oncogene KIT and the chromatin modifiers CHD7 and SUDS3, as well as high expression of genes involved in membrane fusion and budding, and the unfolded protein response.

Among the five TP53-mutated clusters, cluster 4, which has the best survival outcomes, has a loss on chromosome 15, as well as low expression of genes involved in DNA repair and oxidative phosphorylation. Interestingly, tumors in this cluster are enriched for splice-site mutations in TP53 (20% of TP53 mutations in this cluster), and mutations in exon 4 of TP53 (25%), whereas the other clusters are dominated by missense and nonsense mutations in exons 5-10. However, neither exon 4 nor splice site mutations, nor both combined, were significantly associated with survival in this dataset.

Cluster 6 is a small cluster of only 14 samples, associated with high DNA methylation, KRAS mutations and increased expression of the chromatin remodeling factor SATB2. Finally, cluster 8 shows the worst overall survival; it is associated with males, a high rate of point mutations, and low methylation. In addition to TP53 point mutations, it has a loss of 19p (MAP2K7, STK11; >50% samples also have reduced expression of both these genes), high expression of the RNA methyltransferase NSUN2, and high expression of genes for the mitotic spindle, Myc targets, E2F targets, and mTORC1 signaling.

### Head and neck squamous cell carcinoma

Head and neck squamous cell carcinomas are very heterogeneous in aetiology and phenotype. They are stratified by tumor site, stage and histology, and HPV (human papilloma virus) has been associated with better patient outcomes^25^.

We find 8 subtypes of HNSCCs (Figure 3a), which are significantly associated with overall and disease-specific survival. Tumors in clusters 1 and 2 are predominantly HPV+ and TP53 wild-type. They are found mostly in the tonsils and base of tongue, and share a loss on 11q. These HPV+ clusters have significantly higher overall (p=6.1 × 10^-3^) and disease-specific (p=2.1 × 10^-3^) survival than the remaining clusters (3-8).

**Figure 3.**
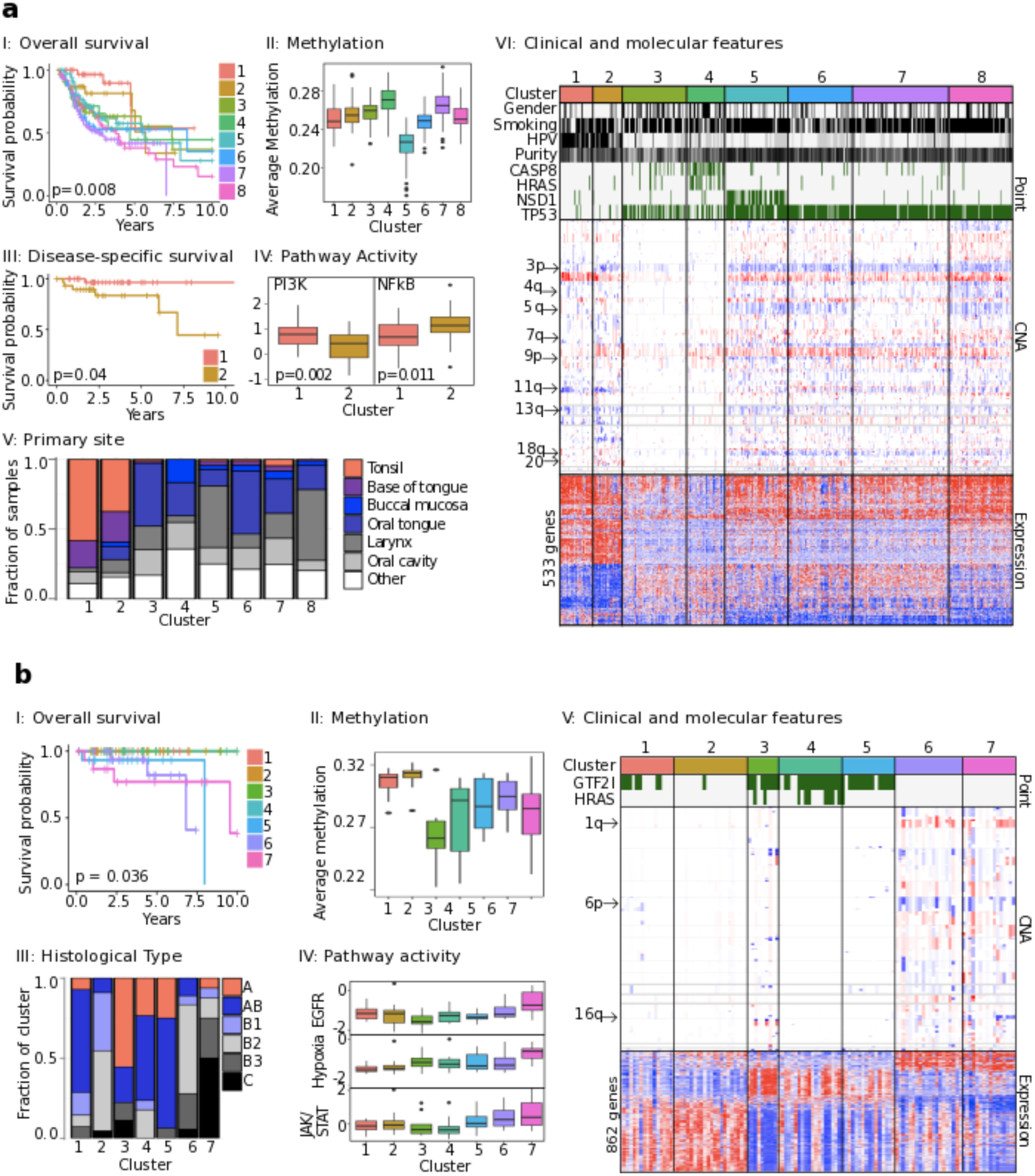
**(a)** Subtyping of 495 head and neck squamous cell carcinomas. I: Overall survival probability for the 8 clusters. II: Boxplot showing average promoter methylation for each cluster. III: Disease-specific survival for clusters 1 and 2. IV: Boxplot showing PROGENy pathway activities for PI3K and NFkB pathways in clusters 1 and 2. V: Bar chart showing the fraction of tumors in each cluster according to primary site of the tumor. VI: Selected clinical and molecular features that differentiate the 8 clusters. For gender, smoking and HPV, gray bars represent missing data. Black bars represent females, smokers, and HPV infection. **(b)** Subtyping of 116 thymomas. I: Overall survival probability for the 7 clusters. II: Boxplot showing average promoter methylation in patients belonging to each cluster. III: Distribution of histological types within each cluster. IV: Boxplot showing pathway activity calculated by PROGENy for EGFR, hypoxia and JAK/STAT pathways, for each cluster. V: Selected clinical and molecular features that differentiate the 7 clusters.

However, cluster 2 has significantly (p=0.041) worse disease-specific survival than cluster 1, and differs in gene expression. While cluster 1 is associated with high expression of 59 genes including the oncogenes DEK and PIK3CA, cluster 2 shows elevated NFKB2 expression, and reduced expression of CDH1 and MAP2K4. GSEA shows that tumors in cluster 2 also show reduced expression of genes involved in PI3K/AKT/mTOR signaling. Consistent with these features, PROGENy shows that cluster 2 has significantly higher NFkB pathway activity than cluster 1, whereas cluster 1 has significantly higher activity of the PI3K pathway. Finally, 62% of the samples in cluster 2 have loss of chromosome 3p, compared to 27% of cluster 1. 3p loss, when occurring jointly with HPV status, has been associated with worse prognosis^26^.

The remaining 6 clusters are HPV-negative and tend to have point mutations in TP53. We do not find any significant survival differences within this group, although they differ in molecular features. Cluster 4 has high DNA methylation and is enriched for females and nonsmokers. This cluster lacks the common 3q gain but is enriched for point mutations in CASP8, FAT1, HRAS, HUWE1 and the histone methyltransferase KMT2B. Clusters 5, 6, 7 and 8 all have high genomic instability. However, cluster 5 is associated with loss of function of the histone methyltransferase NSD1; 68% of the samples have a point mutation in NSD1 while an additional 6% have homozygous deletion of this gene. Tumors in this cluster are hypomethylated, a pattern previously associated with NSD1 loss^27^, and have losses on 13q and 9p.

Cluster 8, which has the highest genomic instability, is enriched for a gain on 7q (including SMURF1; also increased in expression) and a loss on 4q, and high expression of 35 genes including PIK3CA and the transcriptional regulator YEATS2, as well as low expression of the ubiquitin-conjugating enzyme UBE2D3, a phenotype linked to cell cycle progression, reduced apoptosis, and telomere stability^28^. In addition, these tumors show reduced expression of genes for protein secretion, unfolded protein response, and RNA degradation.

### Thymoma

Thymomas are normally classified by histology; however, we found no significant difference in survival between histological types in our data. Instead, CIMLR finds 7 clusters (Figure 3b) with a significant difference in overall survival, each containing a mix of histological types.

Clusters 1 and 2 have high DNA methylation and few mutations or copy number alterations. Cluster 2 is associated with high expression of Myc and E2F targets as well as genes for RNA metabolism, telomere maintenance and DNA synthesis, and low expression of genes for nucleotide excision repair, proteasome and p53 signaling. Clusters 3, 4, and 5 are associated with point mutations in the transcription factor GTF2I, which controls cellular proliferation and has been associated with indolent thymomas^29^.

Clusters 6 and 7 have significantly worse overall (p=2.2 × 10^-3^) and disease-specific (p=2.9 × 10^-3^) survival than the rest of the thymomas. Patients in cluster 6 have a gain on 1q (65% samples) including cancer-associated genes SMYD3, PYGO2, ADAM15, UBE2Q1 and HAX1 (all of these also show increased expression), as well as genes involved in steroid metabolism and phospholipid biosynthesis. 65% also have a loss on 6p including several genes involved in chromatin organization.

Cluster 7 is a mix of histological types, but contains 8 of the 11 type C tumors in the dataset. These tumors share the 1q gain seen in cluster 6; however, only 50% of samples share the 6p loss. In addition, 50% have a loss on 16q, including the tumor suppressor CYLD, several genes for DNA repair (POLR2C, TK2) and chromatin organization (BRD7, CHMP1A, CTCF). POLR2C, TK2, BRD7 and CTCF also show reduced expression in the same samples. This cluster is also associated with increased expression of genes for glycolysis and mTORC1 signaling.

### Clear cell renal cell carcinomas

Clear cell renal cell carcinomas are the most common kidney cancers. Common genetic alterations include mutations in VHL and PBRM1, 3p loss and 5q gain. CIMLR finds two cluster number peaks, at 4 and 10. We first present the results for 4 clusters and then highlight important subclusters found on examining the split into 10 (Figure 4a).

**Figure 4.**
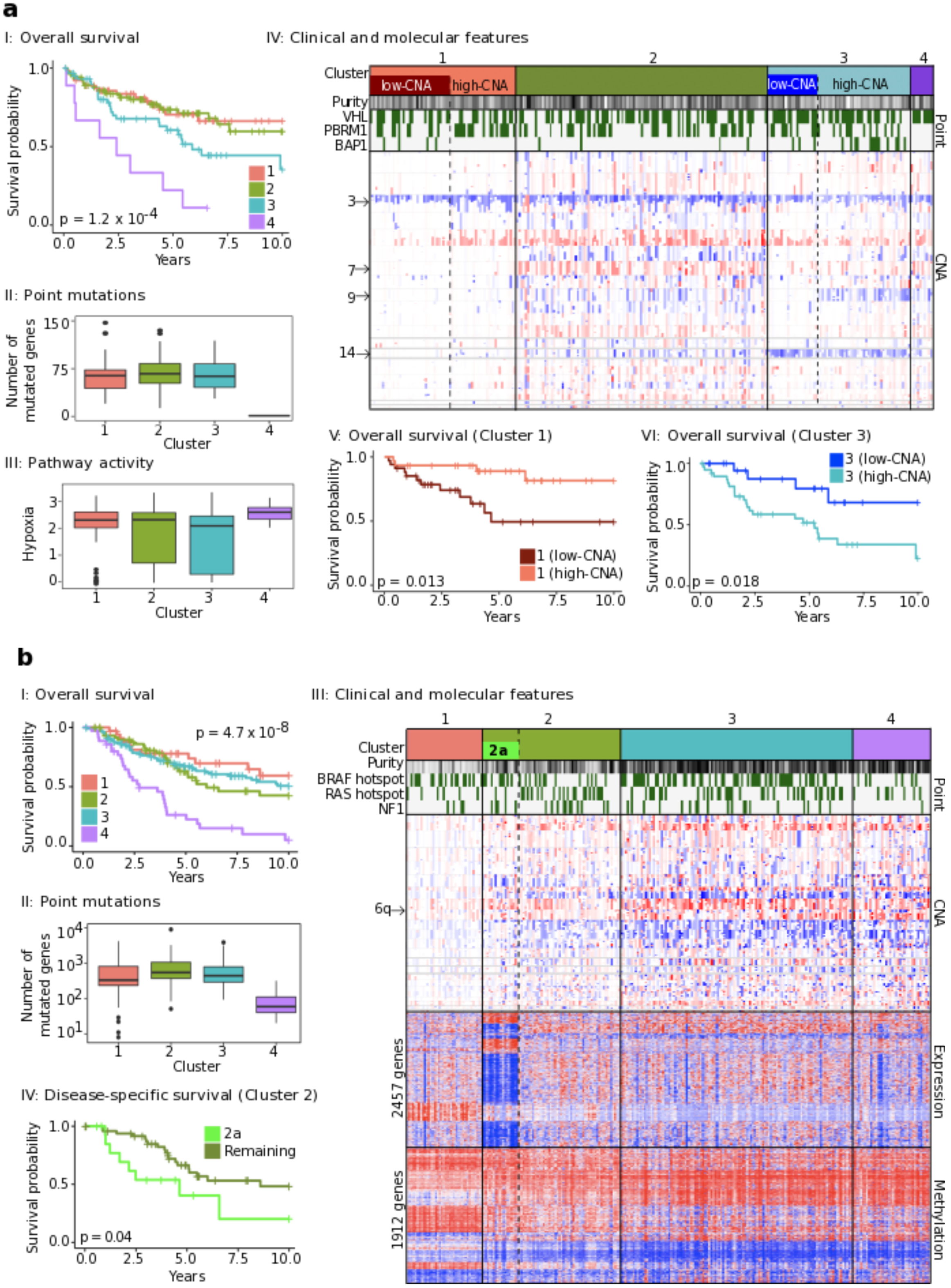
**(a)** Subtyping of 260 clear cell renal cell carcinomas. I: Overall survival probability for the 4 clusters. II: Boxplot showing the number of mutated genes in patients belonging to each cluster. III: Boxplot showing PROGENy pathway activity for hypoxia in the 4 clusters. IV: Selected clinical and molecular features that differentiate the 4 clusters. V: Difference in survival between subsets of cluster 1. V: Difference in survival between subsets of cluster 3. **(b)** Subtyping of 291 skin cutaneous melanomas. I: Overall survival for the 4 clusters. II: Boxplot of the number of mutated genes in patients belonging to the 4 clusters. III: Selected clinical and molecular features that differentiate the 4 clusters. IV: Difference in disease-free survival between 2a and the other clusters.

The clusters show significant differences in overall and disease-specific survival. Clusters 1 and 2 have the best survival outcomes; cluster 2 shows higher genomic instability, particularly a gain on chromosome 7. Cluster 3 has significantly worse survival outcomes than clusters 1 and 2 (p=0.022), and is characterized by a loss on chromosome 14, including the tumor suppressor WDR20, which also shows reduced expression; this gene suppresses growth and apoptosis in renal cancer cell lines^30^.

Finally, cluster 4 is a small cluster of tumors that has significantly worse overall and disease-specific survival than all the other three clusters. These tumors have only one point mutation each in coding regions (mostly in VHL), low expression of the chromatin modifier SETD2, and high expression of the helicase DDX11, which is overexpressed in multiple cancers and associated with proliferation and survival in melanomas^31^.

On examining the split into 10 clusters, we found that several of these smaller clusters were subsets of the 4 major clusters. Interestingly, a subset of cluster 1, characterized by fewer copy number alterations, shows significantly worse overall (p=0.013) and disease-specific (p=0.013) survival than the rest of cluster 1. We also identified a subcluster within cluster 3 which shows significantly better overall (p=0.017) and disease-specific (p=0.004) survival than the rest of cluster 3. This low-CNA group lacks a loss on chromosome 9 (including NOTCH1 and the tumor suppressor TSC1) which is present in the rest of the cluster. Instead, it has reduced expression of several genes involved in DNA repair (CCNK, MLH3, MTA1, APEX1).

### Cutaneous melanoma

Cutaneous melanoma is particularly difficult to subtype since it frequently has a high mutational burden. These tumors have been classified on the basis of common mutations (BRAF hotspot, RAS hotspot, NF1, and triple-WT); however, this classification system is not predictive of patient outcomes^10^. Instead, CIMLR finds 4 clusters significantly associated with overall and disease-specific survival (Figure 4b), and a second-best split at 10. Clusters 1, 2, and 3 are not significantly different from each other in terms of survival outcomes; however, cluster 4 has significantly worse overall and disease-specific survival than all other clusters.

Cluster 1 is characterized by relatively low purity and high immune cell infiltration. Cluster 2 has higher purity as well as high expression of genes involved in mTORC1 signaling and DNA synthesis. While the outcomes for these patients are similar to those for cluster 1, on examining the split into 10 clusters, we identify a subcluster (2a) that has significantly worse disease-specific survival than the rest of cluster 2, and is in fact comparable to cluster 4. This subcluster has a distinctive expression pattern which does not appear to be driven by copy number. This includes high expression of genes for autophagy, organelle fusion and protein transport, and low expression of many genes involved in the G2/M checkpoint, splicing, DNA repair, RNA metabolism, and chromatin remodeling.

Cluster 3 is differentiated from the other clusters by a loss of 47 genes on chromosome 6q and by reduced expression of genes involved in oxidative phosphorylation.

Finally, cluster 4 is distinguished by a much lower point mutation burden (~80 coding mutations per tumor) than all the other clusters, as well as high expression of 3 genes (BTBD9, CDYL, TFAP2A) and high methylation at 100 genes.

### Breast cancer

Breast cancers are frequently classified by intrinsic subtypes^1^ or by the presence of ER, PR and HER2 receptors. Another classification, IntClust^32^, comprises 10 clusters based on selected copy number and expression features. CIMLR obtains 13 clusters (Figure 5a), which are significantly different in overall and disease-specific survival. 10 of these are predominantly ER+ while 3 are predominantly triple-negative. There are significant differences in survival within each group and we examine them separately.

**Figure 5.**
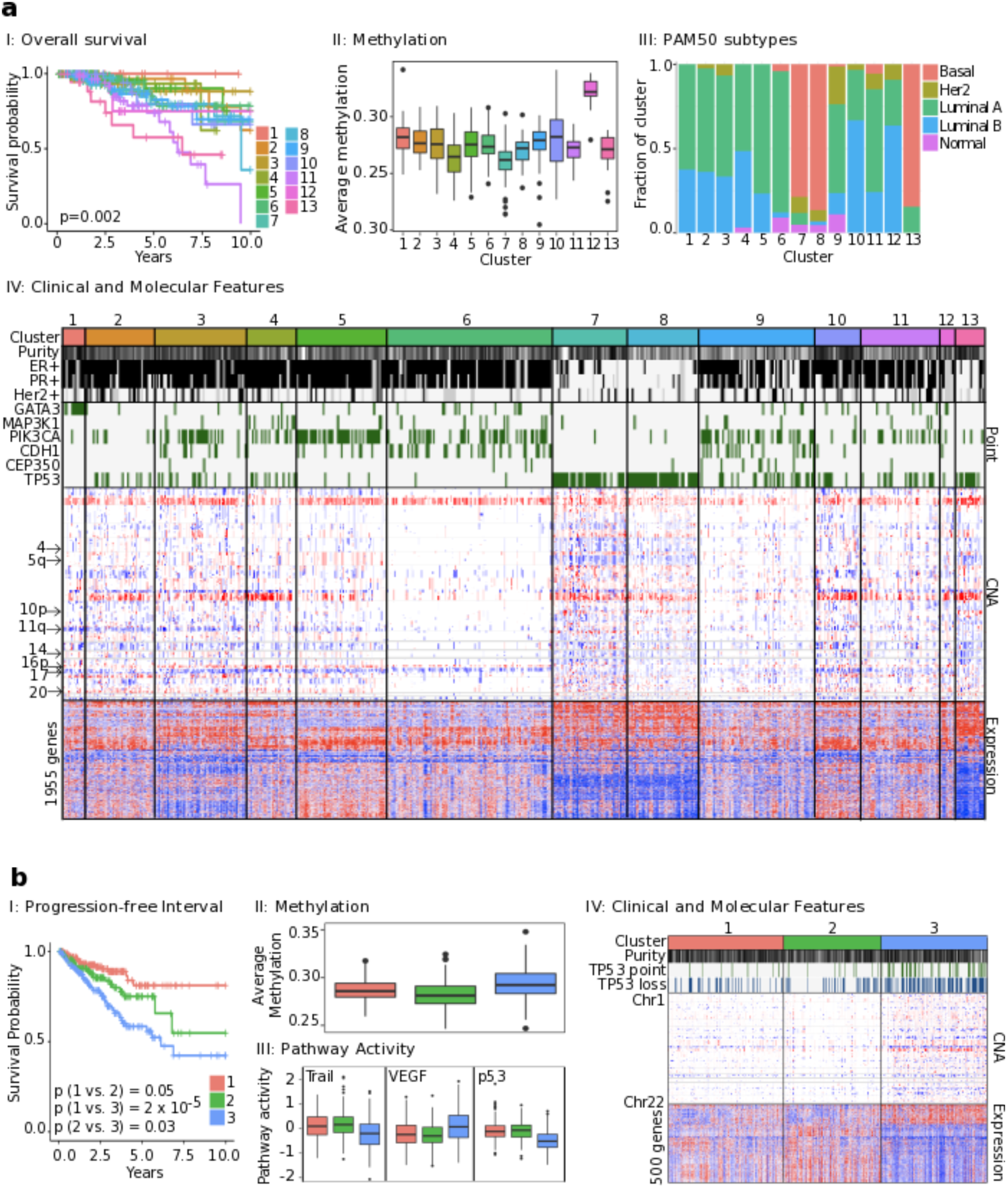
**(a)** Subtyping of 663 breast cancers. I: Overall survival probability for the 13 clusters. II: Boxplot showing average promoter methylation in patients belonging to each cluster. III: Bar plot showing distribution of *PAM50* subtypes within each cluster. IV: Selected clinical and molecular features differentiating the 13 clusters. For ER+, PR+, and HER2+, gray bars represent missing data. **(b)** Subtyping of 490 primary prostate tumors. I: Progression-free intervals for the 7 clusters. II: Average promoter methylation. III: Differences between pathway activities for the 3 clusters. IV: Selected genomic features that differ between the 3 clusters.

Among the predominantly ER+ clusters, Clusters 1, 2, and 3 share a loss on 11q that includes SDHD, ATM, ARHGEF12 and EI24. Cluster 1 has the best survival outcomes and is enriched for point mutations in GATA3 (71% samples). On the other hand, clusters 2 and 3 are enriched for HER2+ tumors and have gains on 17q and 20, as well as a loss on 17p that includes the ssDNA-stabilizing protein RPA1. In addition, Cluster 3 has a gain on 16p, which is shared by clusters 4 and 5, and is enriched for patients with african ancestry. Cluster 12 is a small cluster of 11 patients that display global DNA hypermethylation and high expression of genes involved in telomere maintenance.

Cluster 11 has significantly worse survival outcomes than all the other predominantly ER+ clusters (except clusters 4 and 12 which have small sample size). This cluster is differentiated from the other ER+ clusters primarily by methylation; it shows significant hypermethylation of 128 genes and hypomethylation of 186 genes. It has low expression of NEURL4; this gene encodes a regulator of centrosome organization and its depletion results in mitotic abnormalities in human cell lines^33^. This cluster also has high expression of TAF2, encoding a transcriptional regulator associated with dedifferentiation and proliferation in ovarian cancer^34^. Finally, PROGENy results show that this cluster has significantly higher activity of the MAPK pathway than the rest of the ER+ clusters.

Three clusters - 7, 8, and 13 - are dominated by triple-negative tumors. All three are characterized by TP53 mutations, as well as losses on chromosomes 4, 5q, 15q, and 14q, and a gain on 10p. They also display similar patterns of expression and methylation, and clusters 8 and 13 are enriched for patients with african ancestry. However, cluster 13 has significantly worse survival outcomes than clusters 7 and 8 (p=0.031 and p=0.045 respectively for disease-specific survival). This cluster is differentiated from clusters 7 and 8 by elevated expression of 230 genes and reduced expression of 442 genes including the tumor suppressors APC, CREB1, NCOR1, and NUP98. In addition, it has significantly higher VEGF activity than clusters 7 and 8 according to PROGENy, suggesting higher angiogenesis. It is notable that the 6 ER+ tumors in this cluster share the expression changes described above, suggesting that they may represent a class of aggressive triple-negative-like ER+ tumors.

### Prostate cancer

CIMLR finds 3 clusters in a dataset of 490 primary prostate tumors. Primary prostate tumors have previously been classified on the basis of gene fusions and common point mutations. For this cancer, we do not consider overall survival as very few patients died during the 10-year follow-up period; instead, we observe that the three clusters are significantly different in both progression-free interval (Figure 5b) and disease-free interval.

Clusters 1 and 2 differ primarily in expression, as well as methylation of 13 promoters. Cluster 3 has significantly worse outcomes than both clusters 1 and 2. It is characterized by high genomic instability, including loss of the tumor suppressor TRIM35 on chromosome 8, reduced expression of the tumor suppressor RHOBTB2, and high promoter methylation. It also has higher activity of the VEGF pathway and lower activity of the apoptotic Trail pathway. 63% of the samples in this cluster have p53 mutation and/or loss, and the cluster has lower p53 pathway activity than the others according to PROGENy.

### CIMLR validation on unseen data

We used additional multi-omic datasets to validate our biological findings for the cancers discussed above. For five cancers, we were able to find a sufficient number of samples and data types for validation. For four of these cancers (lower-grade glioma, clear cell renal cell carcinoma, cutaneous melanoma and breast cancer) we were able to obtain new unseen samples recently released by TCGA^35^. For prostate cancer, we used a non-TCGA multi-omic dataset^36^. For each of these cancers, we classified the tumors in the new dataset into risk groups based on the original clusters, using features that differed significantly between clusters (see Methods). We then assessed whether the survival differences discovered in the original dataset were reproduced in the test data (Table 2).

**Table 2:** Validation of CIMLR findings on unseen data. p-values and CI were calculated for overall survival, except for prostate cancer, in which case Disease-Free Interval (DFI) was used. p-values given are for the log-rank test.

| Cancer type | Comparison | Samples (training) | p-value (training) | CI (training) | Samples (test) | p-value (test) | CI (test) |
| --- | --- | --- | --- | --- | --- | --- | --- |
| Lower-grade gliomas | Cluster 1 vs. rest | 282 | $2.0 \times 10^{-25}$ | 0.76 | 226 | $1.8 \times 10^{-12}$ | 0.75 |
| Lower-grade gliomas | Cluster 2a vs. rest of cluster 2 | 147 | 0.035 | 0.63 | 108 | 0.028 | 0.69 |
| Clear cell renal cell carcinoma | (cluster 3 vs. clusters 1+2) | 260 | 0.022 | 0.58 | 138 | 0.019 | 0.63 |
| Cutaneous melanoma | (cluster 4 vs. rest) | 262 | $2.3 \times 10^{-9}$ | 0.59 | 103 | 0.036 | 0.59 |
| Breast cancer (excluding triple-negative) | (cluster 11 vs. rest) | 536 | $3.2 \times 10^{-6}$ | 0.59 | 347 | $1.6 \times 10^{-5}$ | 0.59 |
| Prostate cancer | (Cluster 3 vs. rest) | 490 | $1.2 \times 10^{-3}$ | 0.65 | 118 | 0.011 | 0.62 |

For lower-grade gliomas, we classified 226 tumors into the three major clusters found by CIMLR and validated that Cluster 1 (IDH-wt) has lower survival than the rest of the population. We then selected the 108 tumors predicted to belong to cluster 2 (IDH mutant non-codel) and classified them into high-risk (subcluster 2a) and low-risk (subclusters 2b+2c) groups. Our novel finding that tumors of subcluster 2a have worse overall survival outcomes than the rest of this cluster was validated in this dataset.

For clear cell renal cell carcinoma, we classified 138 samples into high-risk (cluster 4), intermediate risk (cluster 3) or low-risk (cluster 1 + cluster 2) classes. Only 2 samples in the validation set were classified into the high-risk group (cluster 4). However, we showed that samples classified into intermediate risk (cluster 3) had significantly worse overall survival than samples classified as low-risk. For cutaneous melanoma, we classified 103 tumors into high-risk (cluster 4) or low-risk (other clusters) groups. Tumors classified into cluster 4 had significantly worse overall survival.

For breast cancer, we focused on non-triple negative tumors. We classified our validation set of 347 tumors into high-risk (cluster 11) and low-risk (other clusters) groups and showed that tumors classified into cluster 11 had significantly worse overall survival, as was observed in the training set. Finally, for prostate cancer, we classified 118 samples into high-risk (cluster 3) or low-risk groups (cluster 1 + cluster 2) and showed that the samples classified as high-risk had significantly worse outcomes in terms of disease-free interval, which is also consistent with our findings in the training set. This demonstrates that the survival differences discovered by CIMLR are reproducible and potentially clinically useful.

### Prognostic value of CIMLR clusters

In order to ask whether multi-omic subtyping results in prognostic value beyond clinical variables commonly employed to predict survival, we also evaluated the prognostic value of the clusters using Cox proportional hazard regression. For each of the 23 TCGA cancers for which the clusters discovered by CIMLR showed significant association with survival by the log-rank test, we calculated the hazard ratio and associated p-values for each cluster, as well as the Concordance Index (CI)^37^ associated with the clusters. We also calculated these statistics for standard clinical variables provided by TCGA, such as patient age, gender, race, ethnicity, tumor stage and grade, and found that in several cancers (e.g. pleural mesothelioma, cutaneous melanomas, head and neck squamous cell carcinomas) the CI of CIMLR clusters exceeded that of all the tested clinical variables. We then constructed a multivariable Cox regression model for each cancer, including the CIMLR clusters as well as all the clinical variables that were significantly (P<0.1) associated with survival in single-variable Cox regression. In 11 cancers, we found that CIMLR clusters were associated with significant hazard even after adjusting for all tested significant clinical variables (Supplementary Table 10). We also performed the same analysis in the 5 datasets used for external validation of our results. For all comparisons, the CI was similar in the training and test datasets (Table 2). Moreover, for lower-grade gliomas, cutaneous melanomas, and breast cancer, the stratification of the unseen patients on the basis of CIMLR clusters was significantly associated with survival after adjusting for clinical variables (Supplementary Table 11). These results provide strong evidence that multi-omic subtyping using CIMLR offers significant prognostic value beyond that of commonly used clinical features.

## Discussion

The importance of integrative cancer subtyping has been recognized for several years, and multiple algorithms have been developed to exploit the growing amount of available multidimensional data^4,5,6,7,8^. CIMLR addresses many of the weaknesses of current integrative subtyping algorithms, outperforming all tested methods in terms of cluster separation and stability. Furthermore, most of the alternative algorithms proved impractically time-consuming and computationally intensive to run on the considerable volume of data analyzed in this study. As the amount of genomic data is growing at an increasing rate and more types of data are becoming available (such as gene fusions, structural variants, RPPA, miRNA, and ATAC-Seq), efficient methods are essential. Of the available methods, CIMLR is not only superior in terms of performance but is also capable of practically scaling to large-scale analyses with many more data types. We therefore anticipate significant use of this method in the future.

The subtyping achieved by CIMLR demonstrates both biological and clinical relevance. The discovered clusters exhibit significant differences in the activity of oncogenic and tumor suppressor pathways, and they also show significant differences in patient survival in 27 of 36 cancer types. We demonstrate the value of multi-omic subtyping with CIMLR by detailed analysis of 9 cancers. The discovered subtypes provide valuable biological insights and are more predictive of survival than other commonly used classifications. For example, for thymomas the CIMLR subtypes perform better at predicting survival than the histological classifications, while the four CIMLR subtypes of cutaneous melanoma are much better at predicting survival than the earlier mutational classification based on BRAF, RAS and NF1 mutations.

For head and neck squamous cell carcinomas, we separate HPV+ tumors into two groups with significantly different survival outcomes and pathway activity; a previous attempt to subtype these tumors using gene expression did not predict survival^38^. Similarly, in clear cell renal carcinomas, where chromosome 14 loss has been associated with poor prognosis^39^, we not only find a cluster enriched for chromosome 14 loss but show that this is divided into two subclusters only one of which is associated with poor prognosis. This is distinguished by other molecular features including co-deletion of chromosome 9. In primary prostate cancer, we find a pattern of high methylation, copy number alterations, and RHOBTB2 loss that is associated with poor prognosis.

In breast cancer, we identify a cluster of primarily ER+ tumors that have significantly worse survival outcomes than other ER+ clusters; these are distinguished by both methylation and expression features. We also separate the aggressive triple-negative cancers for the first time into three clusters, one of which is considerably more aggressive than the other two and is associated with reduced expression of several well-known tumor suppressor genes. We also find several ER+ and HER2+ samples clustering along with triple-negative cancers and displaying similar expression and methylation patterns.

For five cancers, we validated the significant survival differences discovered by CIMLR in external datasets, showing that CIMLR is capable of discovering molecular subtypes associated with robust, reproducible clinical outcomes.

Our results demonstrate the value of machine learning-based multi-omic clustering in cancer, and the need for more effective and practically usable algorithms. We provide a method for this purpose and anticipate its use in many applications. For example, we expect that subtyping will be useful in stratifying patients for prediction of outcomes and drug response to improve personalized treatment. In addition, our work can be used as a resource for future studies aimed at understanding the biology and evolution of these cancers. As more data becomes available, we expect that the predictive power of subtyping by CIMLR and related approaches will continue to increase and that the medical community will begin to embrace these approaches to improve patient outcomes.

## Methods

### Data preprocessing

We considered all the 32 cancer types studied by TCGA and collected, for each of them, multi-omic data comprising somatic point mutations (as TCGA Mutation Annotation Format files and converted to binary values, 0 to report absence of a mutation in a gene and 1 to report its presence), copy number alterations (log2 ratios between tumor and normal tissue), methylation (Illumina 450; beta-values, i.e. continuous values between 0 and 1) and expression (z-scores normalized to normal tissue or to tumors with diploid genomes). For the TARGET data, we considered 4 pediatric tumors: acute myeloid leukemia, Wilms tumor, neuroblastoma and osteosarcoma. For each of them we collected multi-omic data comprising copy number alterations (log2 ratios between tumor and normal tissue), methylation and RNA expression.

Moreover, we removed extreme values for both copy number log2 ratios and expression z-scores by setting values greater than 10 to 10 and values lower than −10 to −10. We refer to TCGA guidelines for a detailed description of the data obtained from the consortium at the following Website: https://wiki.nci.nih.gov/display/TCGA. All the considered data were within the Open Access Data Tier.

Each data type was modeled as a matrix *N* ×*M*, where N represents the samples, i.e., the patients, and M a set of genes. Each data matrix was normalized so that values ranged between 0 and 1.

### CIMLR

We extended the original implementation of SIMLR^11^ to use multi-omic data. The version of SIMLR adopted here is the default version rather than the large-scale version which leaves out the similarity enhancement by diffusion step.

The original method^11^ takes as input a dataset where rows are samples and columns are genes, and constructs a set of Gaussian kernels for the dataset by fitting multiple hyperparameters. Gaussian kernels are defined as follows: 
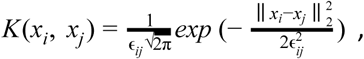
Where *x_i_* and *x_j_* denote the i-th and j-th rows (i.e., samples) of the input data and ϵ^2^_*ij*_ is the variance.

For CIMLR, we represented each of the data types as a patient x gene matrix. We then performed the above procedure for each data type independently, to obtain a set of 55 gaussian kernels with different variance per data type. The number of 55 kernels per data type was empirically derived. Specifically, we assessed the variability of the resulting clusters in terms of normalized mutual information for lower-grade gliomas for a variable number of kernels per data type; 55 kernels represent the point where a plateau is reached (see Supplementary Table 9).

Then, we solved the same optimization problem described in SIMLR^11^, but considering the Gaussian kernels for all the data types together to build one patient x patient similarity matrix. This optimization problem is defined as follows: 
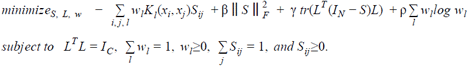

Here, N is the number of patients, C is the number of clusters, i is the row (sample) index, j is the column (gene) index, and l is the kernel index which ranges from 1 to 55 x (number of data types). In the optimization framework, we solve for *S*, i.e., the *N*×*N* similarities matrix; moreover, *w_l_* represents the weight of each Gaussian kernel, *I_N_* and *I_C_* are *N* ×*N* and *C*×*C* identity matrices, β and γ are non-negative tuning parameters, ∥ *S* ∥_*F*_ is the Frobenius norm of *S* and *L* an auxiliary low-dimensional matrix enforcing the low rank constraint on *S*.

### Number of clusters

We also extended the method to estimate the best number of clusters presented in SIMLR^11^ based on separation cost to multi-omics. For a given value of C, we aim at finding an indication matrix Z(R) = XR, with X being the matrix of the top eigenvectors of the similarity Laplacian and R a rotation matrix. Let [*M*(*R*)]_*i*_ =max_*j*_[*Z*(*R*)_*i*,*j*_]. Then, we can define the following cost function to be minimized: 
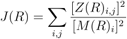

The best number of clusters is the one for which we obtain the largest drop in the value of J(R) over the set of values we consider for C.

We then considered 2 to 15 clusters for the cancer types where we had at least 150 samples, or a maximum of *N* /10 clusters (where N is the number of samples) for smaller datasets.

Cluster assignments for all samples in the study are given in Supplementary Tables 4 and 8.

### Survival analysis

We used four outcome metrics provided by TCGA: Overall Survival (OS), Disease-Specific Survival (DSS), Progression Free Interval (PFI) and Disease Free Interval (DFI), over a time interval of 10 years. For Overall Survival (OS), we censored data points corresponding to patients who died within 30 days or were over the age of 80 at the beginning of the observation period. For TARGET, we only considered Overall Survival (OS) data in the same way as for TCGA. Clusters with only 1 sample were removed prior to survival analysis. Associations between subtypes and outcome were then calculated by Kaplan-Meier analysis using a log-rank test.

Cox regression analysis was performed to estimate hazard ratios associated with individual clusters and to test whether significant associations between clusters and survival outcomes remained after adjusting for common clinical features. Univariate Cox regression was used to select significant (Wald test p<0.1) clinical features which were then included along with CIMLR clusters in a multivariate Cox regression model. Patient age, gender, race, ethnicity, tumor stage and grade were taken into account where data was available. For prostate cancer, Gleason score was taken into account. 5 cancers with an insufficient number of events to fit the Cox regression model were excluded from this analysis.

Survival analysis was carried out using the survival 2.41-3 R package.

### Significant Feature Selection

Molecular features significantly enriched in each cluster were selected as follows. For each cluster, we carried out a hypergeometric test for enrichment of point mutations in each gene. We selected point mutations with an FDR-adjusted p-value of less than 0.05.

To select genes significantly enriched for copy number alterations, we obtained GISTIC thresholded copy number data for each sample from TCGA. We considered a value >=1 to represent gain of the gene and <= −1 to be loss of the gene. For each cluster, we used a hypergeometric test to assess whether the cluster was significantly enriched for either loss or gain of the gene, and selected genes with an FDR-adjusted p-value less than 0.05. For additional stringency and to select the features that were most representative of an individual cluster, we further selected only those genes that were altered in at least 2/3 of the samples in the cluster and <1/3 of the samples in at least one other cluster.

To select expression changes that were significantly enriched within a cluster, we considered a gene to be overexpressed when the z-score was >=1, and underexpressed if the z-score was <= −1. For each cluster, we selected enriched genes using the same criteria as for copy number.

For methylation, we considered a gene to be highly methylated when the beta-value was >=0.75 and unmethylated when the beta-value was <=0.25. For each cluster, we selected genes enriched for high or low methylation using the same criteria as for copy number.

### Classification of unseen data

To classify previously unseen samples into the CIMLR clusters, we used random forest classifiers. Features were ranked on the basis of the hypergeometric test described above and the threshold for selecting the most significant features was tuned to obtain high (>80%) out-of-bag classification accuracy on the training set. We used the ranger version 0.9.0 and caret version 6.0-79 R packages to train random forests and classify unseen samples. For all cancers other than prostate cancer, all 4 input data types were used for classification. For prostate cancer, only expression and copy number data were available for the test set.

### Pathway Analysis and Immune cell infiltration

Gene Set Enrichment Analysis was performed on each cluster using the method of Segal et al.^41^. Gene sets (GO, Cancer Hallmarks, KEGG, Reactome) were obtained from mSigDB^42^. PROGENy pathway activity scores for 11 signaling pathways in TCGA patients were obtained from Schubert et al^17^. Estimates of tumor immune infiltration were obtained from Li et al^19^. All statistical analyses were carried out in R version 3.3.3.

## Author contributions

S.B., B.W., and D.R. designed CIMLR based on SIMLR. B.W. and D.R. implemented the software in MATLAB. D.R. and A.L. processed TCGA data and analyzed the results. A.L. performed cluster annotation, pathway analysis, and external validation. A.L., D.R. and A.S. designed the overall study and drafted the manuscript. All authors read and approved the final manuscript.

## Acknowledgments

We thank Dr. Noah Spies for useful discussions. This work was supported by an R01 grant to A.S. and S.B. (NIH/NCI) and gift funding from the BRCA Foundation. A.L. is supported by a Young Investigator Award from the BRCA Foundation. The results published here are based in part upon data generated by the TCGA Research Network (http://cancergenome.nih.gov/) and by the Therapeutically Applicable Research to Generate Effective Treatments (TARGET) initiative, phs000218, managed by the NCI. Information about TARGET can be found at http://ocg.cancer.gov/programs/target.

## Data Availability Statement

The authors confirm that all relevant data generated in this study are included in the article and/or its supplementary information files.

## Code Availability

The Matlab and R implementations of CIMLR are available at https://github.com/danro9685/CIMLR.

